# Inter-individual Variability of Functional Connectivity in Awake and Anesthetized Rhesus Monkeys

**DOI:** 10.1101/531293

**Authors:** Ting Xu, Darrick Sturgeon, Julian S.B. Ramirez, Seán Froudist-Walsh, Daniel S. Margulies, Charlie E. Schroeder, Damien A. Fair, Michael P. Milham

## Abstract

**Background:** Nonhuman primate models (NHP) are commonly used to advance our understanding of brain function and organization. However, to date, they have offered few insights into individual differences among NHPs. In large part, this is due to the logistical challenges of NHP research, which limit most studies to five subjects or fewer.

**Methods:** We leveraged the availability of a large-scale open NHP imaging resource to provide an initial examination of individual differences in the functional organization of the nonhuman primate brain. Specifically, we selected one awake fMRI dataset (Newcastle: n = 10) and two anesthetized fMRI data sets (Oxford: n = 19; UC-Davis: n = 19) to examine individual differences in functional connectivity characteristics across the cortex, as well as potential state dependencies.

**Results:** We noted significant individual variations of functional connectivity across the macaque cortex. Similar to the findings in human, during the awake state, the primary sensory and motor cortices showed lower variability than the high-order association regions. This variability pattern was significantly correlated with T1w/T2w map, the degree of long-distance connectivity, but not short-distance connectivity. However, the inter-individual variability under anesthesia exhibited a very distinct pattern, with lower variability in medial frontal cortex, precuneus and somatomotor regions and higher variability in the lateral ventral frontal and insular cortices.

**Conclusions:** This work has implications for our understanding of the evolutionary origins of individual variation in the human brain, as well as methodological implications that must be considered in any pursuit to study individual variation in NHP models.

## INTRODUCTION

Our understanding of the human brain has been greatly advanced by continuous interactions between scientists studying the human brain, and those using animal models, which allow for more precise, invasive interrogation of neural circuits. In many cases, our knowledge of the rodent and non-human primate (NHP) is more advanced than that of the human due to the vast array of tools available. However, with rare exception (1), one aspect in which human neuroscience research has outpaced animal research is in the study of individual differences in brain structure and function, and how such differences relate to behavior.

Spurred on by the increased popularity of connectomics research, individual differences in functional connectivity profiles have gained particular attention in the human literature (2–6). A heterogenous picture has emerged in full brain studies, with inter-individual variations being lower in early sensory and motor cortices, and greater in heteromodal association regions, which are characterized by greater long-range connectivity and commonly implicated in higher order cognitive function (2, 7). Putting the pieces together, several have suggested that the increases in functional variation from the primary sensory-motor to unimodal to the heteromodal cortex in humans appear to follow the hierarchy of cortical processing, which has been related to gradients of gene-expression profiles as well (8–10). Importantly, recent studies examining the ratio of T1w to T2w structural MRI images and its links to the cortical hierarchy, have suggested a high degree of similarity between the two species (8, 11). As such, one might expect a high degree of correspondence in patterns of inter-individual variation.

The present work attempts to take a first step in addressing the critical gap in knowledge regarding individual differences in brain organization in NHP. Specifically, we ask the following three questions: 1) How does inter-individual variation in functional connectivity vary throughout the macaque cortex? 2) How does the pattern of inter-individual variation in functional connectivity relate to the connectivity distance, T1w/T2w, and the variation of structural properties? 3) How does the pattern of inter-individual variation in functional connectivity differ between awake and anesthetized states? This latter question is a particularly important one in bridging the gap between the study of individual differences between humans and NHPs, as macaque imaging data is commonly collected in the anesthetized state. This is understandable as it is often more tolerable for the monkey and allows for longer scan times and higher quality data. Nonetheless, differences in functional connectivity between awake and anesthetized states have been described (12–15), raising the possibility that patterns of individual variation may also differ as a function of scan state.

To accomplish our goals, we leveraged previously collected fMRI datasets made available through the PRIMatE Data Exchange (PRIME-DE) (16). Specifically, we used three independent data collections (Oxford [n=20; anesthetized fMRI]; UC-Davis [n=19; anesthetized fMRI]; Newcastle [n = 14; awake fMRI]). To facilitate comparison with findings from the existing human literature, which can provide novel insights into the evolutionary roots of inter-individual differences, our examination followed the framework for evaluation of individual variation previously established by Mueller et al. (2). Specifically, we: 1) examined the spatial distribution of inter-subject variability in functional connectivity while controlling for intra-subject variance, technique noise, and cortical structural profiles, 2) compared inter-subject variation maps to those for cortical structural variation (cortical thickness, curvature, sulcal depth, surface area) and T1w/T2w, and 3) compared inter-subject variation maps to short-range and long-range functional connectivity.

None of the three datasets included in PRIME-DE contained both awake and anesthetized data in the same subjects. Nonetheless, we attempted to provide initial insights into the potential impact of anesthesia. We accomplished this by identifying findings obtained using the awake dataset (Newcastle) that differed from both of the anesthetized datasets (Oxford, UC-Davis). Additionally, we took advantage of the availability of a small sample of macaques scanned both awake and anesthetized (NKI) to corroborate findings suggested by the comparison of data across sites differing with respect to anesthesia state.

## METHODS

We made use of the MRI data from the recently formed NHP data sharing consortium – the non-human PRIMate Data-Exchange (PRIME-DE, http://fcon1000.projects.nitrc.org/indi/indiPRIME.html (16)). Three cohorts of macaque monkeys have been included in the present study.

### Oxford data (anesthetized)

The full data set consisted of 20 rhesus macaque monkeys (macaca mulatta) scanned on a 3T scanner with 4-channel coil. The data were collected while the animals were under anesthesia without contrast-agent. Nineteen macaques were included in the present study (all males, age=4.01±0.98, weight=6.61±2.04); each has a 53.33min (1600 volumes) scan. One macaque was excluded due to the failure of surface reconstruction of its structural image.

### UC-Davis data (anesthetized)

The full data set consisted of 19 rhesus macaque monkeys (macaca mulatta) scanned on a Siemens Skyra 3T with 4-channel clamshell coil. All the animals were scanned under anesthesia without contrast-agent. Nineteen macaques were included in the present study (all female, age=20.38±0.93, weight=9.70±1.58). One resting-state scan (6.67min, 250 volumes) was included per animal.

### Newcastle data (awake)

The full data set consisted of 14 rhesus macaque monkeys (macaca mulatta) scanned on a Vertical Bruker 4.7T primate dedicated scanner. We restricted our analysis to 10 animals (8 males, age=8.28±2.33, weight=11.76±3.38) for whom two awake resting-state fMRI scans were required. The details of the data acquisition and experimental procedures were described in previous studies (17–23). No contrast-agent was used during the scans.

### NKI data (awake and anesthetized)

The NKI data consisted of 2 rhesus monkeys (macaca mulatta) scanned on a 3.0 Tesla Siemens Tim Trio scanner with an 8-channel surface coil. This dataset was not used for variation analysis but as a complementary dataset, which consists of a large amount of awake and anesthetized data within the same monkey. The data had been preprocessed and used in our previous work (13). Specifically, we included 7 sessions (awake: 224 min, anesthetized: 196.7 min) for monkey-W, and 4 sessions (awake: 86 min, anesthetized: 108 min) for monkey-R. All the included data from NKI site were collected with contrast agent monocrystalline iron oxide ferumoxytol (MION). The details of the acquisition parameters were described in Xu et al. (13).

### MRI Data Preprocessing for NHP datasets

The structural MRI data were processed using the customized HCP-like pipeline from the Fair’s lab and the functional data were preprocessed as described previously (13, 24). See details of the preprocessing in Supplemental Method.

### Human data (awake) and processing

We replicated prior human finding here by using resting-state fMRI data from Human Connectome Project (HCP) dataset (177 unrelated participants, 79 male, age=29.03±3.45) (24, 25). The minimal preprocessing HCP data were temporal filtered (0.01-0.1Hz), smoothed (FWHM=6mm) and down-sampled to 10k mid-cortical surface.

### Functional connectivity Profile (FC profile)

In order to account for the intra-subject variance while estimating the inter-subject variability, we first split the data into several subsets for each animal. Specifically, for Oxford data, we split the entire session into 5 subsets (320 volumes per subset). For UC-Davis data, which has less time points, we split the data into 2 subsets (125 volumes per subset). For the Newcastle data, two scans were originally recorded per animal (250 volumes per scan). In order to limit the intra-subject variation within the scan, we also split each scan into 2 subsets (125 volumes x 2 subsets x 2 scans). For Human HCP data, we kept the original 4 resting-state fMRI scans as 4 subsets (1200 volumes per subset).

Similar to earlier human studies (26), we first defined the functional connectivity profiles at each vertex as a vector of Pearson’s correlations between its time course and the time courses of all the regions of interest (ROIs). The ROIs were uniformly sampled from a coarse sphere surface (624 vertices per hemisphere) and mapped back onto the 10k sphere in Yerkes19 macaque space (27); each ROIs includes one resampled vertex and its one-step neighboring vertices while the ROIs that fell in the medial wall were removed from the analysis (Figure 1B). In total, there were 1,114 ROIs for two hemispheres. For a given vertex *v*, let the connectivity profile denote as a vector *FC_v_*(*subject, subset*). *FC_v_* is a 1 × 1114 vector representing the functional connectivity for a given vertex v to 1,114 ROIs. A similar procedure was employed to the human HCP data to create a connectivity profile (a 1 × 1114 FC vector) at each vertex v.

**Figure 1.**
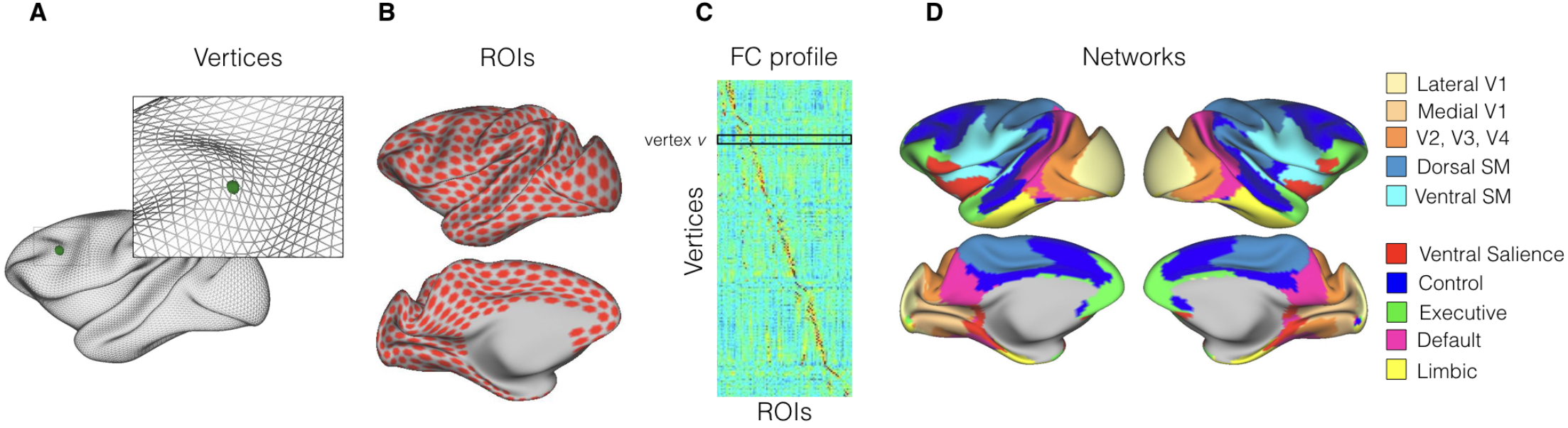
The functional connectivity profile and the functional networks of macaque brain. (A) The inflated macaque surface mesh (10,242 vertices). (B) The uniformly sampled ROIs on the 10k macaque surface (1,114 ROIs within the cortex mask for two hemispheres). (C) The example of the functional connectivity profile. Each row is a vector of functional connectivity profile calculated as the correlation between the time series at vertex v and 1,114 ROIs. (D) The large-scale functional networks based on the clustering approach.

### Inter-subject Variability of the FC profile

Following the same analysis framework in prior studies (2), the distance of functional profile *FC_v_* between any subsets and subjects was calculated by 1 - Pearson’s correlation at each vertex. The intra-subject variability was quantified by averaging the distance of *FC_v_*(*subject,subset*) across subjects as follows:

*V_v_*(*intra*) = 1 − *E*(*corr*[*FC_v_*(*subject,i*), *FC_v_*(*subject,j*)]), where *i* and *j* indicates scanning subsets for the same subject.

The inter-subject variability was initially estimated as:

*V_v_*(*inter_unadjusted*) = 1 − *E*(*corr*[*FC_v_*(*p,subset*),*FC_v_*(*q,subset*)]), where *p* and *q* indicate different subjects.

Of note, the inter-subject variability was adjusted by regressing out the intra-subject variability, the technical noise (temporal signal-to-noise, tSNR), and the group-averaged cortical structural profile using the general linear model (GLM). Four commonly examined cortical measures were included in the model (i.e., thickness, sulcus depth, surface area and curvature). The adjusted inter-subject variability was estimated as follows:

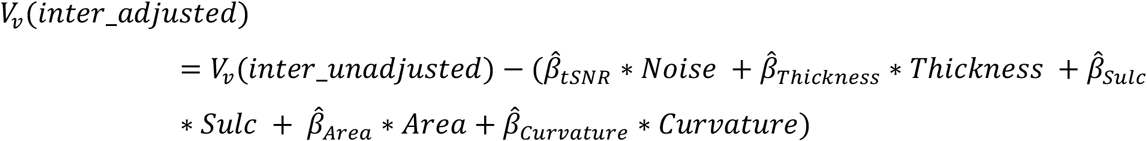

### Inter-subject Variability of Cortical Structure

We also measured the inter-subject variability for the cortical structural measures (i.e. curvature, sulcus depth, surface area, cortical thickness) by calculating the standard variation within each site and scaling the averaged measurement and its square term in GLM model at each vertex *v* as follows:

*Group_Std_v_*(*X*) = 1 + *β*_1_* *Group_Mean_v_*(*X*) + *β*_2_* ((*Group _Mean_v_*(*X*))^2^, where X represents a given cortical measure.

### Removing the Geometrical Artificial Effects

Of note, when mapping the data from the volume to the surface and smoothing the data on the surface, the connectivity between surface vertices with a certain range of geodesic distance might be artificially increased. To remove such artificial connectivity, we explicitly removed the local connectivity (the geodesic distance between vertex *v* and ROIs < 6mm on the surface) from the *FC_v_*(*subject, subset*) vector for for each vertex. To demonstrate this artifactual effect of intrinsic cortical geometry, we performed the randomization test by replacing the real time series with the temporal surrogate data and replicated the above inter-subject variability analysis (Figure S1F-G) without removing the local FC (geo < 6mm). Next, based on the same surrogate data, we tested whether removal of the local FC (geo < 6mm) is effective to eliminate the artifacts. When local connectivity (geo < 6mm) was not removed from the analysis, the randomization results exhibited an artificial spatial distribution of the inter-subject variability which was caused by the cortical folding and the mesh geometric properties (Figure S1F). When we removed the local FC (geo < 6mm) from the analysis, the induced artificial effect was eliminated (Figure S1G). This confirmed the validity and the necessity of removing of the local FC (geo < 6mm) in the analysis.

### Functional Network and Parcellation

To quantify the variation at network level, we performed a clustering approach to identify large scale networks in macaque monkeys. The main goal in the current study is not to delineate optimal clusters but rather to summarize the functional variation in a suitable set of networks which is presumably comparable to the human networks in prior studies (2, 26). Therefore, we conducted the same clustering approach used in Yeo’s study and opted to use the Oxford data which contains the largest sample size and the most extensive data (53.33 min per animal). The details of the clustering method were described in Yeo (26). Briefly, we utilized the functional connectivity profiles based on 1,114 ROIs that sampled uniformly on the surface cross left and right hemispheres. The same clustering algorithm was applied to the group averaged 17,749 × 1,114 (vertices x ROIs) correlation matrix. To determine the number of clusters that is comparable with Yeo’s 7 networks in human, we tested the number of clusters a priori from 7 to 12. The resultant visual and somatomotor networks in macaque were divided into small clusters in a very early stage when 7 networks were chosen. Specifically, the somatomotor network was divided into ventral and dorsal units and been stabilized as 2 clusters from 7 to 12 networks. Similarly, the visual network was divided into lateral V1, V2-V3 and medial V1 clusters and stabilized as 3 clusters from 9 to 12 networks. In Yeo’s 7 networks, 5 networks were identified for heteromodal cortex besides the visual and somatomotor networks. To make the number of heteromodal clusters the same as in humans, ten clusters were chosen for the final demonstration - 5 unimodal networks capture the primary sensory and motor areas (i.e., medial V1, lateral V1, V2-V4, ventral somatomotor, and dorsal somatomotor networks) and 5 heteromodal networks characterize the association cortex (Fig-1D). They resemble the executive network (green), the control network (blue), the ventral salience network (red), the default mode network (magenta), and the limbic network (yellow).

### The Short- and Long-distance FC

To measure the short-distance FC, we thresholded for each vertex by at the top 10% percentile of connectivity strengths and counted the percentage of the connections whose geodesic distance are within 12 mm. Similarly, the long-distance FC was defined as the percentage of the connections whose geodesic distance are greater than 20 mm in the top 10% of all connectivity. As mentioned above, to avoid the geometrical artificial effects, the FC (geo < 6mm) were removed when we calculated the local FC here. Of note, given that the head motion may induce artificial connections, in addition to the scrubbing procedure in preprocessing step, we adopted more strict thresholds and calculated the short- and long-distance FC (Fig-S2) to replicate the analysis.

### Randomization test for correction of spatial correlation

To investigate the spatial correlation of two vertex-wise maps, we proposed the following randomization test to correct the Pearson’ correlation p-value. Specifically, we first generated 96 ROIs (diameter=12mm on the surface) uniformly sampled on the macaque template surface per hemisphere. The distances between any pairs of ROIs are larger than 12 mm on the surface. Then we randomly chose 1 vertex per ROIs to create a sub-sample. This procedure guarantees that the distances between any vertices in the sub-sample is larger than 12 mm, thus the vertices in the sub-sample are most likely to be independent samples, thereby yield a valid p-value from the sub-sample. We then randomly rotated the ROIs along the surface 1000 times to generate 1000 sub-samples and calculate the r and p from each randomization. The correlated r-value was calculated as the average of Pearson’s r across 1000 randomizations. We also counted the proportion of how many randomizations yielded significant p (<0.05) and used 1 minus this proportion for the corrected p-value.

## RESULTS

### Inter-individual Variation of Functional Connectivity

For each data collection selected from PRIME-DE (Newcastle, Oxford, UC-Davis), we estimated inter-individual functional variation at each vertex (see Fig-S1A), while controlling for intra-individual variation, technical noise, and cortical structural properties (i.e., thickness, curvature, sulcus depth, and area) (see Fig-S1B-D). In each of the datasets, inter-subject variability of FC exhibited a heterogeneous spatial distribution across macaque cortex.

We first replicated the prior human findings based on the HCP data (Fig-2A). The heteromodal association cortex particular frontoparietal network showed higher inter-individual variation than the visual and somatomotor networks. Intriguingly, the inter-individual variation was specifically high in the association neworks borders. For the macaque data, focusing on the findings from the data collection observed using the awake state first (i.e., from Newcastle) (Fig-2B), the heteromodal association cortex showed a higher degree of individual differences while the unimodal regions (i.e. visual and somatomotor area) were relatively low. This topographic distribution is consistent with the prior findings in awake human imaging, which showed higher variation in the association cortices and lower variation in visual and somatomotor areas (Fig-2A). To quantify the functional variation at the network level, the vertex-wise variation maps were averaged within each network (Fig-1D). Figure 3 showed the boundary of each network overlaid with the parcel-wise functional variation demonstrated based on parcellation (28). For the awake condition (Fig-3B), the visual, in particular the medial V1 area showed the least individual difference, followed by the lateral V1 and secondary visual area (V2-V4). Among the heteromodal networks, executive network (green) and control network (blue) exhibited the highest variation in FC. The ventral salience network (red) and default mode network (magenta) showed a moderate degree of variation, which is lower than executive and control networks but higher than that of the limbic network across five heteromodal networks. This hierarchy of the functional variation across networks in macaque is consistent with the prior human work (Fig-3A, Mueller et al., (2)) where the highest level of functional variability was observed in frontoparietal control network, followed by the attention, default and limbic networks with a moderate degree of variation; the sensory-motor and visual networks showed the least variation. Intriguingly, unlike the human findings with a steep increase of inter-individual variation from unimodal to heteromodal networks, the variation across networks was relatively mild in macaque (Fig-3A-B).

**Figure 2.**
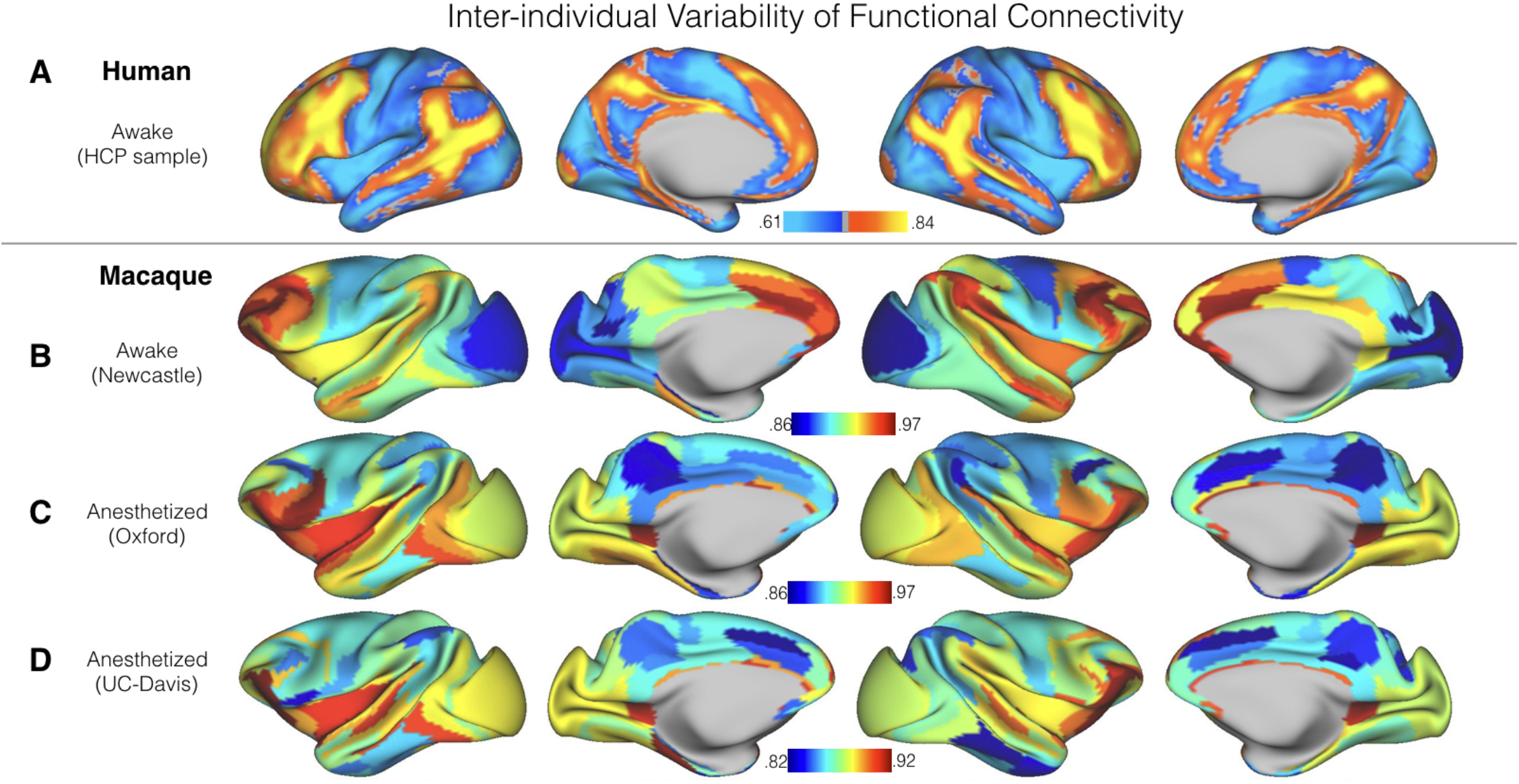
Inter-individual variation of functional connectivity in (A) human HCP sample, (B) awake macaque from Newcastle dataset and (C-D) anesthetized macaque samples (C: Oxford dataset; D: UC-Davis dataset). To emphasize the overall regional differences and improve the SNR for macaque samples, we visualized the inter-individual variation using the architectonic areas from published atlas (28, 57). The vertex-wise maps were demonstrated in Figure S1A.

**Figure 3.**
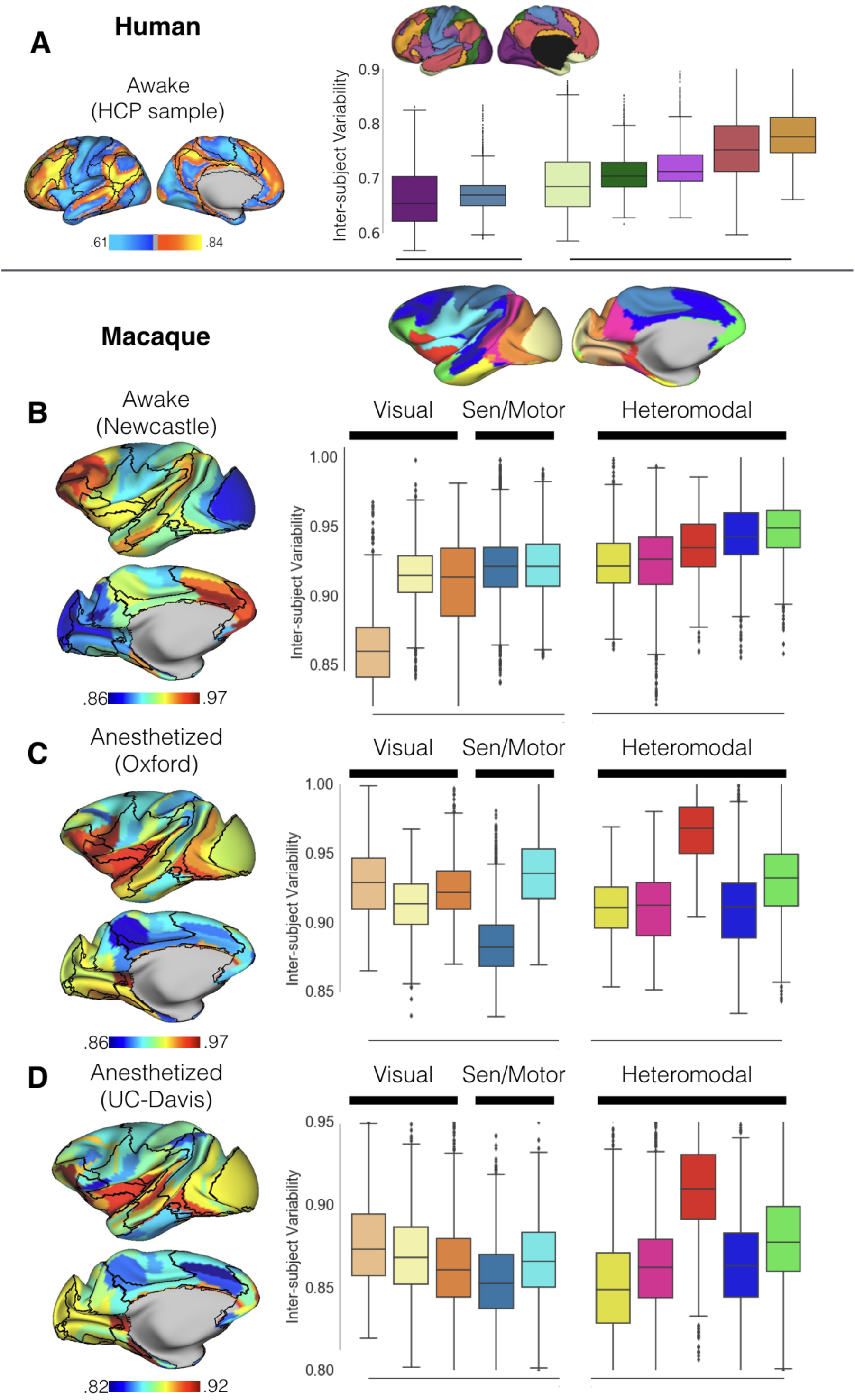
Inter-individual variation of functional connectivity at the network level in (A) awake human HCP sample, (B) awake macaque sample from Newcastle dataset, and (C-D) anesthetized macaque samples (C: Oxford dataset; D: UC-Davis dataset). The boxplot quantified the vertex-wise inter-individual variations using Yeo 7 networks (26) for human sample and the macaque 10 networks (Fig 1D) for macaque sample (B-D).

Next, we looked at the inter-individual variation in each of the collections obtained during the anesthetized state (Fig-2C-D). Notably, the pattern of functional variation observed throughout the cortex was highly similar across the two sites (r=0.43, p<0.001), even despite the relatively smaller amount of data for UC-Davis data (3.3min x 2 subsets per animal) with respect to the Oxford data (10.6 min x 5 subsets per animal). In contrast, findings from the two anesthetized samples appeared to be relatively distinct from those obtained during the awake state at Newcastle, as well as those previously reported from the human. Specifically, individual differences obtained in the anesthetized samples exhibited notably greater functional variation in FC patterns across individuals in visual cortex, and remarkably less in cingulate cortex, precuneus, and area 7m (Fig-2C-D). At the network level, the ventral salience network (i.e. area F5, ProM, Gu, OPRO, and insula) showed the highest degree of variation in the anesthetized state and dorsal somatomotor cortex the lowest, while in the awake samples, the executive function network exhibited the highest degree of variation and medial primary V1 the least.

### Functional Variation and Short- and Long-Distance Connectivity

Next, for each data collection, we assessed the short-range (i.e. local) and long-range (i.e. distant) functional connectivity at each vertex tested for associations with inter-individual variation in functional connectivity. Focusing on the awake sample (Newcastle), long-range FC was most prominent in the lateral and medial frontal cortices, as well as the PCC and inferior temporal cortex (Fig-4A). The spatial pattern of long-range FC showed a strong positive association with the individual variations (r=0.56, corrected p<0.001) across macaque cortex. When we looked at the short-range FC pattern, a non-significant correlation was obtained (r=-0.10, corrected p=0.87) with interindividual variation. The overall patterns of association between inter-individual variation and short-/long-range FC followed those observed in human literature, but were more modest for short-range FC in the macaque (r=-0.10 versus r=-0.33 in Mueller et al. (2).

**Figure 4.**
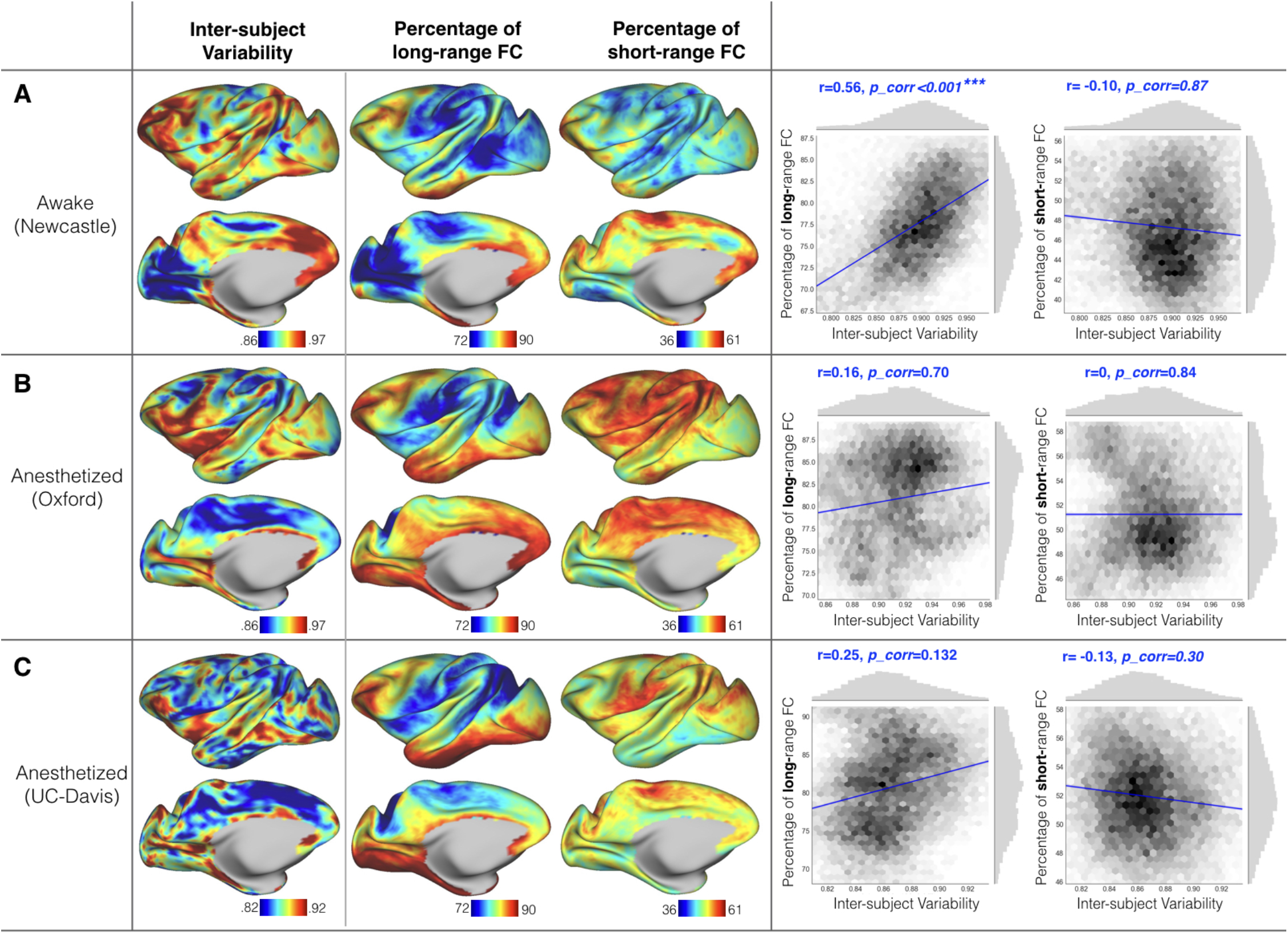
Inter-individual variation of functional connectivity, and the percentage of long-- and short-range connectivity in (A) awake macaque sample from Newcastle dataset, and (B-C) two anesthetized samples (B: Oxford dataset, C: UC-Davis dataset) The vertex-wise maps were shown on the left panel; the scatter plots and the spatial correlations with corrected p-values were shown on the right panel. The randomization test for correcting the p-values were described in Supplemental Method.

In the anesthetized samples (i.e., UC-Davis, Oxford), no similar positive correlations between either long-range or short-range FC and inter-individual variation were found in both Oxford (long-range: r=0.25, corrected p=0.132; short-range: r=0, corrected p=0.84) and UC-Davis (long-range: r=0.16, corrected p=0.7; short-range: r=-0.13, corrected p=0.3) data (Fig-4B-C). Of note, to avoid the geometrical artificial effects as mentioned in the method, the FC that geodesic distance < 6mm was not counted in the short-range FC calculation. This procedure may mitigate the correlation between inter-individual variation and short-range FC.

### Functional Variation and Structural Variation

Human studies have reported associations between inter-individual variations in functional connectivity and structure, particularly with respect to cortical folding pattern (i.e., sulcal depth). Here we investigated the extent to which such correlations may exist in the macaque, focusing on four commonly examined measures (i.e. curvature, sulcus depth, surface area, cortical thickness). At the vertex-level, we did not find any significant correlations between inter-individual variation in functional connectivity and that in the structural indices examined, regardless of sites (all corrected p>0.217).

### Functional Variation and T1w/T2w topography

T1w/T2w mapping has been suggested to index myelin content and at a minimum, appears related to the anatomical hierarchy in both human and macaque (8, 11, 29, 30). Here we assessed the relationship of each, inter-individual variation, short-range functional connectivity and long-range functional connectivity, with T1w/2w measured from a new population average macaque altas referred to as “Yerkes19” (27). A moderate to strong correlation between T1w/T2w and the inter-subject functional variability (r=−0.466, corrected p=0.001) was observed in the awake sample (Fig-5A), though no correlations were found in either of the anesthetized samples (Fig-5B-C). Similarly, we found that T1w/T2w was negatively associated with long-range FC in the awake sample (r=-0.491, corrected p=0.006, Fig-5) while such correlations were not significant in anesthetized sample (Oxford: r=-0.251, corrected p=0.107, UC-Davis: r=-0.163, corrected p=0.555). When correlating the T1w/T2w with the short-distance FC, the awake sample showed a relatively low correlations with a marginal significance (Newcastle: r=-0.20, corrected p=0.058) while no significant correlations were obtained in anesthetized samples (Oxford: r=-0.056, corrected p=0.906; UC-Davis: r=-0.131, corrected p=0.807).

**Figure 5.**
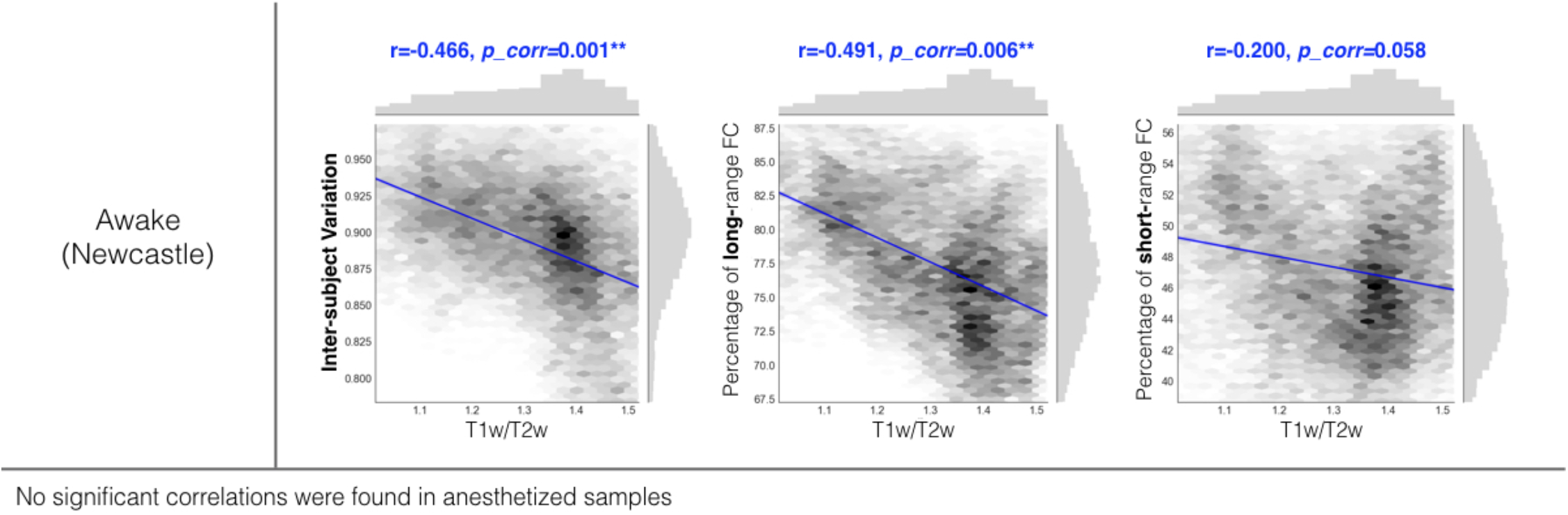
The scatter plots and the spatial correlations between T1w/T2w topography with inter-individual variation of functional connectivity, short- and long-range connectivity in awake macaque sample (Newcastle dataset).

### Quantifying Awake vs. Anesthetized State Differences at the Group-level

Our findings for interindividual variation pointed to the importance of state-related differences. Though it was not a primary focus of the present work, we examined the impact of state on group-level findings as well. To gain further insight into the state-related functional differences, at each vertex, we examined the similarity of functional connectivity profiles derived from awake and anesthetized samples. The similarity was quantified by Pearson correlation coefficient of FC profiles at each vertex (Fig-6). Two specific comparisons were used for the assessment of similarity: 1) the first session from Newcastle (awake) and Oxford (anesthetized) data, and 2) the second session from Newcastle (awake) and UC-Davis (anesthetized) data. Since the site effect confounded with the state differences here, we made such comparisons to test whether the state similarity could be replicable in independent samples (Fig-6A-B). The similarity from these two comparisons showed highly similar patterns (r=0.59, corrected p<0.001). Functional connectivity profiles in unimodal regions (i.e. visual, somatomotor networks) were more similar (reddish color in Fig-6A-B) between awake and anesthetized samples. The heteromodal networks in general showed higher dissimilarity than unimodal regions. Specifically, the limbic (yellow) network exhibited highest dissimilarity, followed by the ventral salience (red), and default mode networks (magenta); the executive network exhibited lowest dissimilarity among heteromodal network between awake and anesthetized samples (Figure-6A-B), though still higher than the unimodal networks.

**Figure 6.**
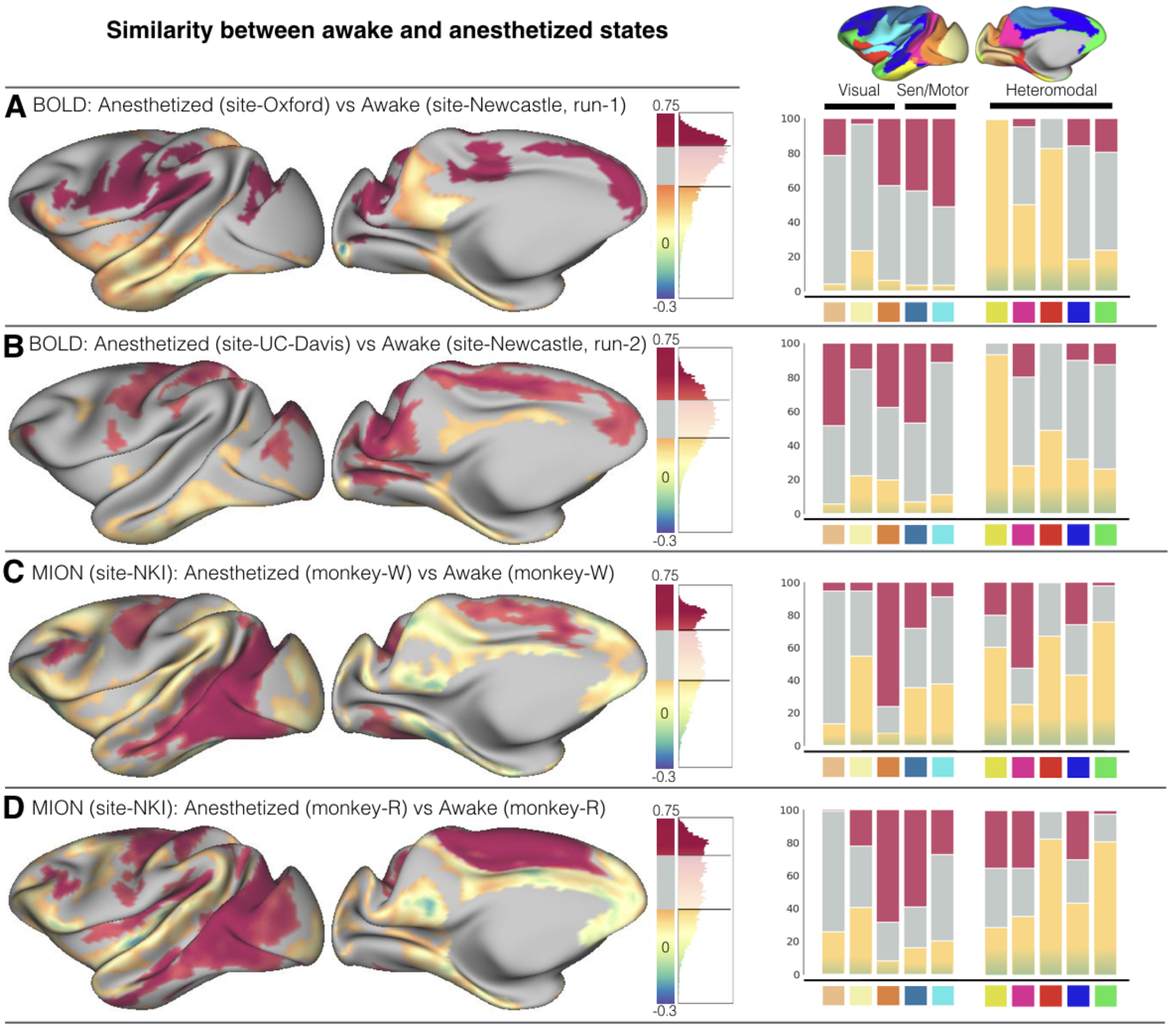
State-related dissimilarity at the group-level. The averaged similarity between awake and anesthetized samples measured as the Pearson correlation of FC profile at each vertex. The distribution of the r-values is visualized beside the surface map for four comparisons: (A) anesthetized sample from Oxford dataset vs. awake sample from the first scan of Newcastle dataset (both Oxford and Newcastle data were BOLD fMRI scans without the contrast agent); (B) anesthetized sample from UC-Davis dataset vs. awake sample from the second scan from Newcastle dataset (both Oxford and Newcastle data were BOLD fMRI scans without contrast agent); (C) anesthetized vs. awake within one macaque from NKI dataset (monkey-W) which were scanned with contrast-agent MION; (D) anesthetized vs. awake within one macaque from NKI dataset (monkey-R) which were scanned with contrast-agent MION. The negative r indicates the opposite pattern of FC profile between awake and anesthetized samples. We mapped the top 25% of positive r-values (the most similar) in reddish and bottom 25% of positive r-values in yellow and all the negative r-values (the most dissimilar) in blueish color. The right panel shows the proportion of most similar (red) and dissimilar vertices (yellow to blueish) lying within each functional network.

To provide further corroborating evidence for state differences, we also quantified state similarity within the same animal using data from two macaque subjects collected at NKI. Though the gross awake vs. anesthetized similarity maps for Monkey-W (Fig-6C) and Monkey-R (Fig-6D) were not significantly related to those for the comparisons of Newcastle versus Oxford/UC-Davis (r=0.11 to 0.23, corrected p=0.533 to 0.063), the heteromodal networks appeared to be more dissimilar relative to unimodal regions (Fig-6). Of note, the NKI awake data were collected under a naturalistic viewing tasks and presumably had visual network activation (13), while the Newcastle awake data were mainly focused on the auditory paradigm (17, 20). In addition, the NKI-data were collected with the contrast agent (MION) which has been demonstrated to be a significant factor that can cause the differences of FC (Supplemental Results), as shown in previous studies (13, 31–33).

## DISCUSSION

Echoing prior reports from human studies (2), the present work suggests that macaque cortex is similarly heterogeneous with respect to individual differences. Consistencies between prior findings in the human and those in the macaque awake imaging sample (Newcastle) examined here were multifold. First, heteromodal association areas were characterized by relatively higher levels of inter-individual variation in functional connectivity patterns, while lower order regions including primary sensory and motor areas were characterized by lower levels of variation in the awake brain. Areas within the lateral prefrontal cortex showing the highest inter-individual variability consistently across the macaque and human (Figure 2A-B). These include area 8B and 9/46v, which, although topographically shifted across species, demonstrated consistent patterns of variability in comparison to neighboring areas. Second, regional differences in individual variation were highly correlated with the presence of long-range functional connectivity and the T1w/T2w map. No significant correlations were observed with the various structural indices, though a modest correlation with sulcal depth variation was observed in humans. Overall, these findings suggest a conservation of patterns of individual differences across species. Of note, our preliminary examination of the impact of anesthesia on individual variation found differences in supplementary motor, insular and lateral inferior frontal cortices, suggesting the potential importance of considering state when attempting to translate imaging findings across states and species.

From an evolutionary perspective, the findings of the present work are informative. The consistency of findings with those of prior human studies suggests that the functional organization of lower order areas (i.e., primary sensory-motor, unimodal areas) were relatively conserved across individuals and heteromodal association cortices were more variable (2, 7, 34). In considering overall patterns of regional differences in the present work, it is worth noting their relations to phylogenetic ordering, with heteromodal association areas emerging later in the evolutionary process (35–38). Leading hypotheses (e.g., the tethering hypothesis) emphasize the contributions of environmental factors (e.g., molecular gradients), and their interactions with genetically-determined processes, to the cortical expansions that led to the development of heteromodal association areas (39). Similar factors may very well contribute to the higher degree of individual differences observed in these areas, which are characterized by a more prolonged developmental course (2, 40).

Both human and macaque awake samples suggested that the unimodal-heteromodal hierarchy was negatively related to an index of anatomical hierarchy - T1w/T2w maps (2, 11, 29, 30). The latter has been suggested as an in vivo measure of myelin content, which has been shown remarkably correlated with the hierarchical gradients in gene expression profiles that related to synaptic physiology, cell-type specificity, and cortical cytoarchitecture (8). Additionally, consistent with the findings in human, the inter-individual variability of FC was positively correlated with the long-distance FC. These long-distance connections are prevalent in hub regions that underlie attention, memory and other higher cognitive functions in humans (2, 41–43). Such strong correlation in macaque and human suggests that long-range FC is critical to the individual differences both within- and cross-species. Taken together, the relationship with connectivity distance, cortical gradients of potential myelin content (i.e. T1w/T2w), and related gene expression profile (8) provides hints at the biological underpinnings of variability of functional connectivity.

While anesthetized imaging is often the most practical approach in nonhuman primate imaging due to the greater ease of handling and tolerability, state-related sensitivities have been reported in both human and nonhuman primates (44–48). The present study was not optimally designed to analyze anesthesia effects, findings do suggest that patterns of individual difference may be impacted by the use of anesthesia. First, in both of the two anesthetized samples employed, we found the limbic and the default networks to be less variable than unimodal regions, while the orbitofrontal cortex and insular cortex were highly variable - the latter of which is commonly linked to the salience network and consciousness in the literature (44, 49, 50). Second, associations between individual variation and long-range connectivity were only detectable in awake sample. These findings suggest that anesthesia may impair our ability to detect variations in long-range functional architecture - a finding that is consistent with prior work that reported decreased network complexity. This may explain observations of decreased variation for posterior portions of the default network.

The high degree of similarity in the findings for the two anesthetized samples was particularly notable given that different anesthesia and imaging protocols were employed. At a minimum, the present work raises cautions for the study of individual differences under the anesthetized state and necessitates future work to clarify anesthesia effects more definitively. This clarification is especially important given prior working emphasizing the presence of changes in the functional architecture associated with sleep and anesthesia, using a range of modalities (e.g., fMRI, EEG, PET, TMS). As NHP imaging researchers move beyond initial demonstrations of similarity in the gross network architecture to more nuanced comparative anatomy questions related to within- and across-species variation, state-related differences will become an increasing concern for NHP studies; if substantive enough, they may necessitate awake imaging depending on the region(s) being examined. An understanding of state-related differences will also be helpful in explaining variations in findings that will likely emerge across studies over time.

There are a number of notable limitations of the present work. First, although larger than what is included in most studies, and comparable to that employed by Mueller et al.(2), the awake sample employed was still relatively modest in size. As such, while we are relatively confident of the gross patterns detected, which well replicated those previously documented in humans, more finely-grained details were likely beyond detectability. Second, the methods and protocols used across sites were determined independently of one another, increasing the likelihood of finding differences among samples. Despite this, a strength of our findings was that the two anesthetized samples yielded highly similar results, and the awake sample replicated the prior human findings. Third, is that the awake and anesthetized comparisons were carried out across samples; ideally, the same subjects would be used for any comparison of states. We did include the two subjects from NKI, which were each scanned under both states, finding corroborating evidence for the between-sample comparison. Finally, while our findings are largely consistent with those previously obtained in the human, we were not able to carry out direct statistical comparisons of findings across species. Future work would benefit from protocol harmonization (as possible) and more rigorous cross-species alignment to provide more fine-grained comparisons (38, 51, 52).

In summary, the present work demonstrates the presence of individual differences in the functional organization of the macaque cortex that are highly reminiscent of what is observed in the human. Similar to the findings in human literature, the present work draws particular attention to differences in the regional properties of the cortical hierarchy, which appear to play out in a growing number of neuroscientific findings (8, 30, 53–56). Finally, it raises cautions about the study of individual differences in the anesthetized state and the need for further attention toward determining which properties are and are not stable across states as scientists work to put together the pieces across studies and species.

## Acknowledgements

This work was supported by gifts from Joseph P. Healey, Phyllis Green, and Randolph Cowen to the Child Mind Institute and grants from the NIH (BRAIN Initiative R01-MH111439 to C.E.S. and M.P.M.; P50MH109429 to C.E.S.; R01-MH107508 to E.L.S. and D.A.F.; and P60-AA010760, R01-MH115357, R01-MH096773, and P50-MH100029 to D.A.F.). This work was also supported in part by NIH grant P01AG026423 and the Yerkes National Primate Research Center base grant (Office of Research Infrastructure Programs; OD P51OD11132). We would also like to thank the investigative teams from Newcastle (J. Nacef, C.I. Petkov, F. Balezeau, T.D. Griffiths, C. Poirier, A. Thiele, M. Ortiz, M. Schmid, D. Hunter), Oxford (J. Sallet, R.B. Mars, M.F.S. Rushworth) and UC-Davis (M. Baxter, P. Croxson, J. Morrison), as well as the funding agencies that made their work possible (UC-Davis: NIA; Newcaste: National Center for 3Rs, NIH, Wellcome Trust, UK Biotechnology Biological Sciences Research Council; Oxford: Wellcome Trust, Royal Society, Medical Research Council, UK Biotechnology Biological Sciences Research Council).

