## Supplemental Figures for "Inter-individual Variability of Functional Connectivity in Awake and Anesthetized Rhesus Monkeys"

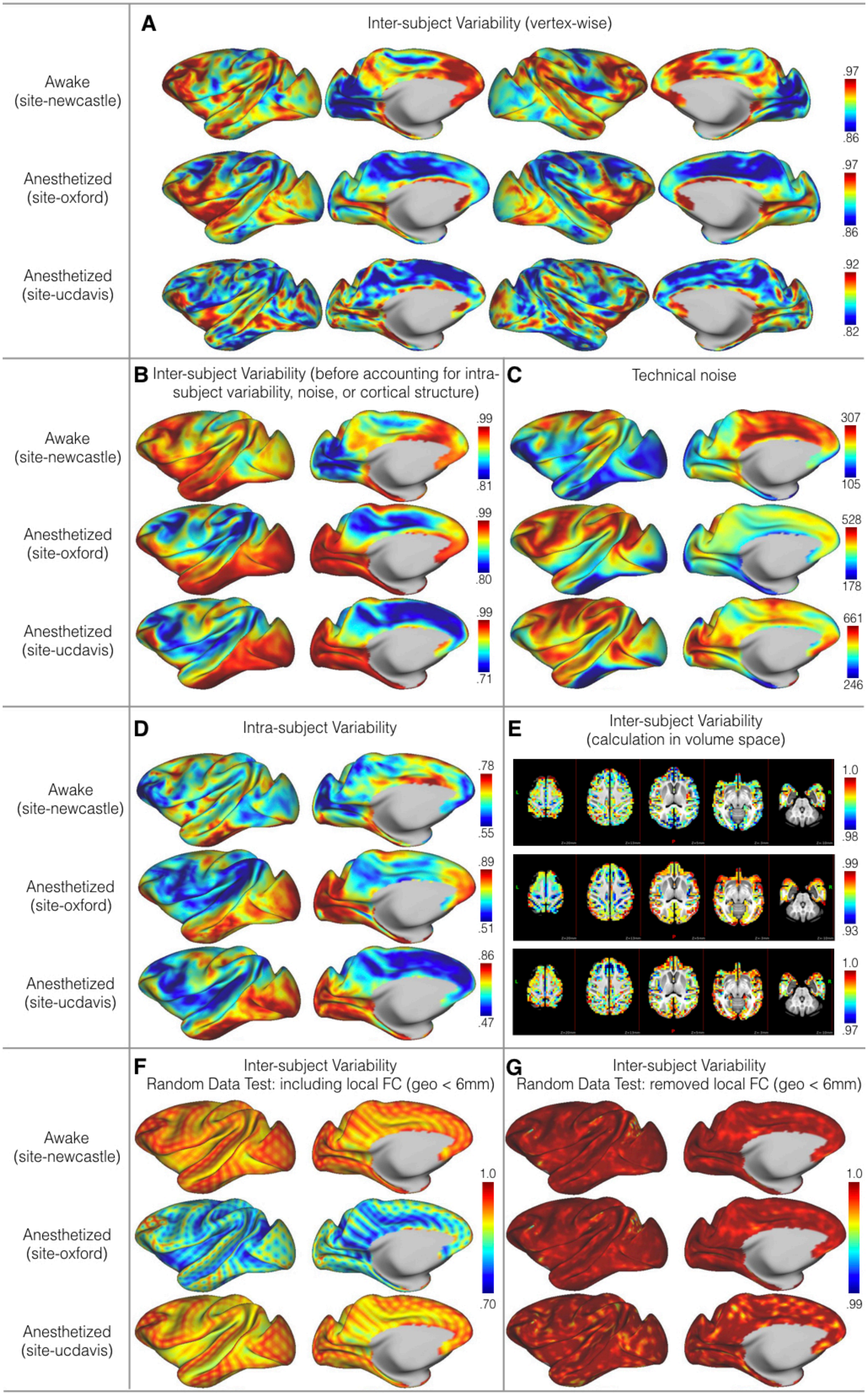
A
Inter-subject Variability (vertex-wise)
Awake (site-newcastle)
Anesthetized (site-oxford)
Anesthetized (site-ucdavis)
.97
.86
.97
.86
.92
.82
B
Inter-subject Variability (before accounting for intra-subject variability, noise, or cortical structure)
.99
.81
.99
.80
.99
.71
C
Technical noise
307
105
528
178
661
246
D
Intra-subject Variability
.78
.55
.89
.51
.86
.47
E
Inter-subject Variability (calculation in volume space)
1.0
.98
.99
.93
1.0
.97
F
Inter-subject Variability
Random Data Test: including local FC (geo < 6mm)
1.0
.70
G
Inter-subject Variability
Random Data Test: removed local FC (geo < 6mm)
1.0
.99

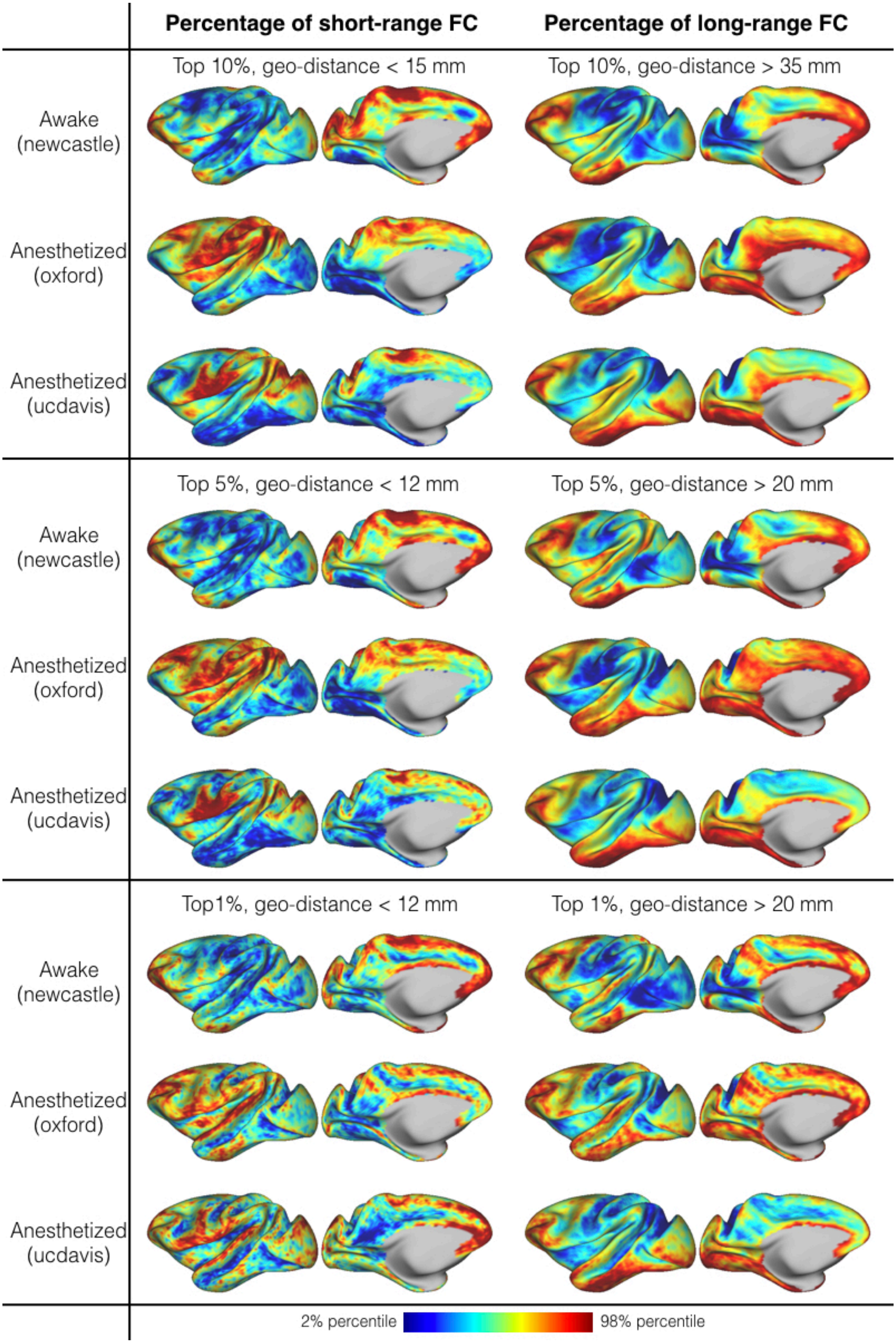

Percentage of short-range FC
Percentage of long-range FC
Top 10%, geo-distance < 15 mm
Top 10%, geo-distance > 35 mm
Awake (newcastle)
Anesthetized (oxford)
Anesthetized (ucdavis)
Top 5%, geo-distance < 12 mm
Top 5%, geo-distance > 20 mm
Awake (newcastle)
Anesthetized (oxford)
Anesthetized (ucdavis)
Top1%, geo-distance < 12 mm
Top 1%, geo-distance > 20 mm
Awake (newcastle)
Anesthetized (oxford)
Anesthetized (ucdavis)
2% percentile
98% percentile

A) The distribution of the variability scores for the matched human samples

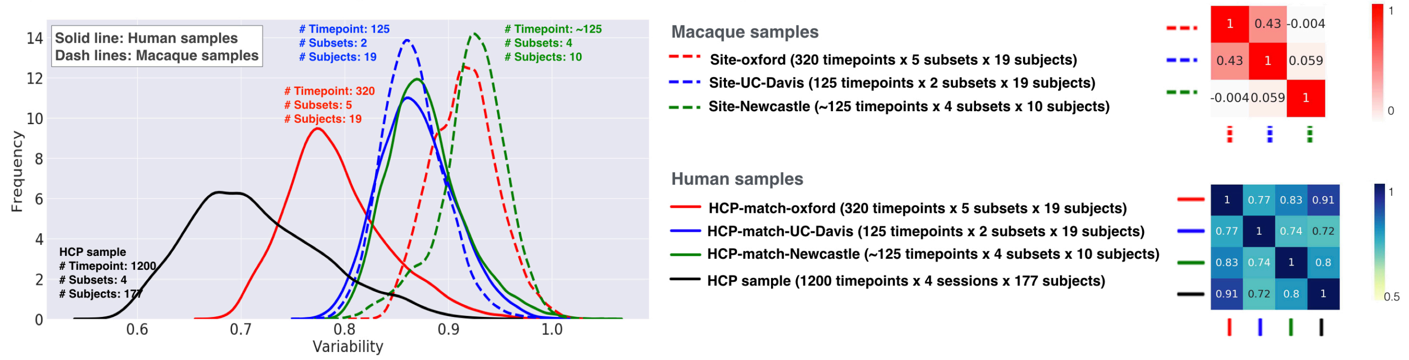


B) The distribution of the variability scores for human sample based on different amount data

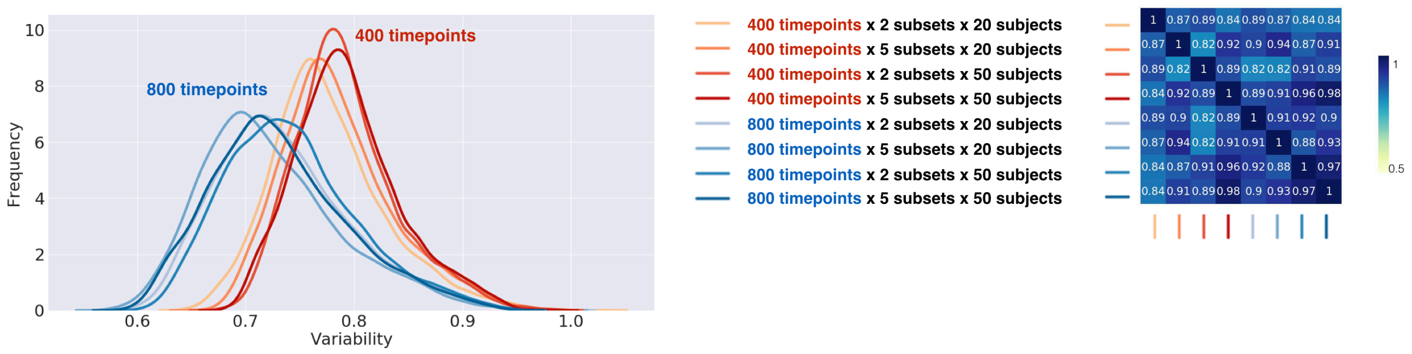
