## Supplemental Methods for "Inter-individual Variability of Functional Connectivity in Awake and Anesthetized Rhesus Monkeys"

**Supplemental Method**

**Macaque Oxford data (anesthetized).** The full data set consisted of 20 rhesus macaque monkeys (macaca mulatta) scanned on a 3T scanner with 4-channel coil. The data were collected while the animals were under anesthesia. Briefly, the macaque was sedated with intramuscular injection of ketamine (10 mg/kg) combined with either xylazine (0.125-0.25 mg/kg) or midazolam (0.1mg/kg) and buprenorphine (0.01 mg/kg). Additionally, macaques received injections of atropine (0.05 mg/kg, i.m.), meloxicam (0.2 mg/kg, i.v.), and ranitidine (0.05 mg/kg, i.v.). The anesthesia was maintained with isoflurane. The details of the scan and anesthesia procedures were described in Noonan et al. (Noonan et al. 2014) and the PRIME-DE website (http://fcon_1000.projects.nitrc.org/indi/PRIME/oxford.html). Resting-state fMRI data were collected with 2x2x2 mm resolution, TE=19ms, TR=2000ms, flip angle=90°. T1-weighted images required using MPRAGE sequence (resolution=0.5x0.5x0.5mm, TE=4.01mx, TR=2500ms, flip angle=8°).

**Macaque UC-Davis data (anesthetized).** The full data set consisted of 19 rhesus macaque monkeys (macaca mulatta) scanned on a Siemens Skyra 3T with 4-channel clamshell coil. All the animals were scanned under anesthesia. In brief, the macaques were sedated with injection of ketamine (10 mg/kg), dexmedetomidine (0.01 mg/kg), and buprenorphine (0.01 mg/kg). The anesthesia was maintained with isoflurane at 1-2%. The details of the scan and anesthesia protocol can be found at (http://fcon_1000.projects.nitrc.org/indi/PRIME/ucdavis.html). The resting-state fMRI data were collected with 1.4x1.4x1.4mm resolution, TE=24ms, TR=1600ms). No contrast-agent was used during the scans.

**Newcastle data (awake).** The full data set consisted of 14 rhesus macaque monkeys (macaca mulatta) scanned on a Vertical Bruker 4.7T primate dedicated scanner. We restricted our analysis to 10 animals (8 males, age=8.28±2.33, weight=11.76±3.38) for whom two awake resting-state fMRI scans were required. The fMRI data were acquired while the animals were awake (resolution=1.2x1.2x1.2mm, TE=16ms, TR=2000ms). Two 8.33-min (250 volumes) scans were acquired for each animal. The structural T1-weighted images were acquired using MDEFT sequence with 0.6x0.6x0.6mm resolution, TE=6ms, TR=750ms. No contrast-agent was used during the scans.

**NKI data (awake and anesthetized).** The NKI data consisted of 2 rhesus monkeys (macaca mulatta) scanned on a 3.0 Tesla Siemens Tim Trio scanner with an 8-channel surface coil. we included 7 sessions (awake: 224 min, anesthetized: 196.7 min) for monkey NKI-W, and 4 sessions (awake: 86 min, anesthetized: 108 min) for monkey NKI-R. All the included data from NKI site were collected with contrast agent monocrystalline iron oxide ferumoxytol (MION, 1- mg/gk iv). The awake scans were acquired with naturalistic viewing (movie watching) paradigm. For the anesthesia scans, the macaques were sedated with an initial does of ketamine (8 mg/kg IM), atropine (0.05 mg/kg IM), and dexdomitor (0.02 mg/kg IM) and maintained anesthesia at 0.75%.

**MRI Data Preprocessing for NHP datasets**. The structural processing includes 1) spatial denoising by a non-local mean filtering operation (Zuo and Xing 2011), 2) brain extraction using ANTs registration with a reference brain mask followed by manually editing to fix the incorrect volume (ITK-SNAP, www.itksnap.org) (Yushkevich et al. 2006); 3) tissue segmentation and surface reconstruction (FreeSurfer) (Dale, Fischl, and Sereno 1999; Fischl et al. 1999); 4) the native white matter and pial surfaces were registered to the Yerkes19 macaque surface template (Donahue et al. 2016). The functional MRI data preprocessing included temporal compressing (3dDespike), motion correction, 4D global scaling, nuisance regression (white matter, cerebrospinal fluid, and Friston-24 motion parameters), band-passed filtering (0.01-0.1Hz), linear and quadratic detrending, boundary-based co-registration from functional space to native anatomical space, projection to native mid-cortical surface created by averaging the white and pial surfaces, and smoothing with 3 mm FWHM along the native mid-cortical surface. The final functional data were then down-sampled to a 10k resolution surface (10,242 vertices per hemisphere) (Donahue et al. 2016).

The Frame-wise displacement (FD) was calculated to quantify the head motion (Power et al. 2012, 2014). In the Oxford and UC-Davis data sets, which were collected under anesthesia, the FDs were less than 0.2 mm for all animal scan volumes. For awake data in the Newcastle dataset, we used 'scrubbing' to remove the volumes with FD>0.2mm. In 16 out of 20 sessions (80% of data in awake sample) less than 20% of time points had to be scrubbed. The final mean FD was 0.066 (SD=0.018) across all awake scans of Newcastle data, 0.019 (SD=0.006) across all anesthetized scans of Oxford data, and 0.028 (SD=0.007) across all anesthetized scans of UC-Davis data.

**Results:**

**The scale of the variability scores.** Noted that the range of the raw variability score in human from the prior studies and the current work (Fig 2A) are lower than the range in monkeys (Fig 2B-D). This is mainly attributable to the non-biological factors (e.g., the dataset sample size, resolution of surface mesh, and data quality). To provide the potential insights, we repeated the analyses by matching the HCP sample specifications (the number of timepoints, sessions and subjects) of each of three NHP samples (Figure S3, left panel). With the same sample specifications, the range of variability scores for matched HCP samples were notably increased. Regardless of the sample specifications, the relative regional difference in inter-individual variation is stable (Figure S3, right panel).

**MION Effect on the Comparison of Awake vs. Anesthetized State Differences.** In Figure 6, the NKI-data were collected with the iron-based contrast agent (MION). Unlike BOLD, fMRI signal with MION are influenced by cerebral blood volume (Leite et al. 2002). MION has been demonstrated to increase the SNR compared with the BOLD signal and commonly used in NHP imaging when scanning at 3.0T scanner (Gautama, Mandic, and Van Hulle 2003; Grayson et al. 2016; Gautama 2003). The MION data were shown similar regional difference of functional connectivity with BOLD fMRI (n.d.). However, it has been demonstrated to be a significant factor that can cause the differences of FC, as shown in previous studies (Gautama, Mandic, and Van Hulle 2003; Grayson et al. 2016; Xu et al. 2018; Leite et al. 2002).

**Figure legend**

**Figure S1.** (A) The vertex-wise inter-individual variability of functional connectivity for three macaque samples. (B) The vertex-wise inter-individual variability of functional connectivity before accounting for intra-individual variability, temporal SNR, or the cortical structure including curvature, thickness, sulcal depth, and surface area. (C) The temporal SNR for three macaque samples. (D) The intra-individual variability of functional connectivity. (E) The inter-individual variability calculated in volume space. (F) The inter-individual variability of random data test without removing the local FC (geo < 6mm). The variation map exhibited an artificial spatial distribution of the inter-subject variability which was caused by the cortical folding and the mesh geometric properties. (G) By removing the local FC from the analysis, the artificial effect was eliminated.

**Figure S2.** The long-range and short-range functional connectivity at different thresholds.

**Figure S3.** The distribution of the inter-individual variability scores in macaque and human samples. (A, left panel) The distribution of individual variability scores for HCP sample, three macaque samples, and three matched HCP samples. The sample specification (# of subject, # of sessions, # of timepoints) affected the scale of variability. (A, right panel) The spatial correlation among samples with different specifications. (B) The distribution of the inter-individual variability scores in human samples. The variability varied with the sample specifications, in particular the time points within each subset.

Gautama, Temujin. 2003. *Motion Processing in Macaque Middle Temporal Area: Computational Principles, Models and Data Analysis*. Leuven University Press.

Gautama, Temujin, Danilo P. Mandic, and Marc M. Van Hulle. 2003. "Signal Nonlinearity in fMRI: A Comparison between BOLD and MION." *IEEE Transactions on Medical Imaging* 22 (5): 636–44.

Grayson, David S., Eliza Bliss-Moreau, Christopher J. Machado, Jeffrey Bennett, Kelly Shen, Kathleen A. Grant, Damien A. Fair, and David G. Amaral. 2016. "The Rhesus Monkey

Connectome Predicts Disrupted Functional Networks Resulting from Pharmacogenetic Inactivation of the Amygdala." *Neuron* 91 (2): 453–66.

Leite, Francisca P., Doris Tsao, Wim Vanduffel, Denis Fize, Yuka Sasaki, Larry L. Wald, Anders M. Dale, et al. 2002. "Repeated fMRI Using Iron Oxide Contrast Agent in Awake, Behaving Macaques at 3 Tesla." *NeuroImage* 16 (2): 283–94.

Noonan, Maryann P., Jerome Sallet, Rogier B. Mars, Franz X. Neubert, Jill X. O'Reilly, Jesper L. Andersson, Anna S. Mitchell, Andrew H. Bell, Karla L. Miller, and Matthew F. S. Rushworth. 2014. "A Neural Circuit Covarying with Social Hierarchy in Macaques." *PLoS Biology* 12 (9): e1001940.

Power, Jonathan D., Kelly A. Barnes, Abraham Z. Snyder, Bradley L. Schlaggar, and Steven E. Petersen. 2012. "Spurious but Systematic Correlations in Functional Connectivity MRI Networks Arise from Subject Motion." *NeuroImage* 59 (3): 2142–54.

Power, Jonathan D., Anish Mitra, Timothy O. Laumann, Abraham Z. Snyder, Bradley L. Schlaggar, and Steven E. Petersen. 2014. "Methods to Detect, Characterize, and Remove Motion Artifact in Resting State fMRI." *NeuroImage* 84 (January): 320–41.

Xu, Ting, Arnaud Falchier, Elinor L. Sullivan, Gary Linn, Julian S. B. Ramirez, Deborah Ross, Eric Feczko, et al. 2018. "Delineating the Macroscale Areal Organization of the Macaque Cortex In Vivo." *Cell Reports* 23 (2): 429–41.

Yushkevich, Paul A., Joseph Piven, Heather Cody Hazlett, Rachel Gimpel Smith, Sean Ho, James C. Gee, and Guido Gerig. 2006. "User-Guided 3D Active Contour Segmentation of Anatomical Structures: Significantly Improved Efficiency and Reliability." *NeuroImage* 31 (3): 1116–28.

Zuo, Xi-Nian, and Xiu-Xia Xing. 2011. "Effects of Non-Local Diffusion on Structural MRI Preprocessing and Default Network Mapping: Statistical Comparisons with Isotropic/anisotropic Diffusion." *PloS One* 6 (10): e26703.

n.d. bioRxiv. https://doi.org/10.1101/155952.
